# Automated profiling of social behaviors to assess the genetic basis of evolution of aggressive behaviors in *A. mexicanus*

**DOI:** 10.1101/2025.07.07.663556

**Authors:** Renee Mapa, Stefan Choy, Mia Lapko, Alli Kimmel, Isabel Carino-Bazan, Johanna E. Kowalko

**Affiliations:** Department of Biological Sciences, Lehigh University, Bethlehem, PA

## Abstract

Across the animal kingdom, social behaviors such as aggression are critical for survival and reproductive success. While there is significant variation in social behaviors within and between species, the genetic mechanisms underlying natural variation in social behaviors are poorly understood. A central challenge to investigating the mechanisms contributing to the evolution of social behaviors is that these behaviors are typically complex, making them a challenge to quantify. The Mexican tetra, *Astyanax mexicanus,* is a powerful model for investigating the evolution of traits, as it is a single species that exists as populations of eyed, river-dwelling surface fish and blind cave-dwelling fish. The blind cavefish have evolved morphological and behavioral differences compared to surface fish, including reduced aggression. Here, we developed and validated an automated machine learning pipeline that integrates pose-estimation and supervised behavioral classification to track and quantify aggression-associated behaviors—striking, following, and circling. Using this pipeline, we established that these behaviors are quantitatively different between surface and cave fish during juvenile stages in *A. mexicanus*, similar to what was observed previously in adults. Moreover, assessment of these aggressive behaviors in surface-cave F2 hybrid fish revealed that striking and following are strongly positively correlated, while striking and circling are negatively correlated, suggesting that these behaviors evolved through some shared genetic mechanisms. These findings demonstrate the power of automated tracking and behavioral phenotyping in multiple fish in *A. mexicanus* and establish a foundation for future studies investigating the genetic basis of evolution of social behaviors.

## Introduction

Aggression, defined as an action that causes harm to another individual, is observed across the animal kingdom (Archer, 1988; Huntingford and Turner, 1987). Across species, aggressive behaviors serve multiple functions to improve reproductive success and survival (Moyer, 1968). For example, *Mus musculus* attack prey such as insects, demonstrating how predatory aggression can aid individuals when securing resources for survival (Butler, 1973). Adult cichlid fish, *Cichlasoma dimerus*, exhibit aggression to protect their offspring and defend spawning sites through behaviors such as biting and chasing (Alonso, Cánepa, Moreira, and Pandolfi, 2011). Aggression has also been shown to be crucial in establishing dominance hierarchies in species such as wild wolves, *Canis lupus,* that exist in hierarchical groups called packs (Sands and Creel, 2004; Schenkel, 1967). The spiny lizard, *Sceloporus jarrovi*, exhibits territorial aggression against predators and conspecifics, particularly during breeding season, to enhance survival and reproductive success (Ruby, 1978). While aggression is a behavior critical for survival and reproduction across multiple species, the genetic mechanisms that contribute to the evolution of aggressive behaviors are poorly understood.

The Mexican tetra, *Astyanax mexicanus,* is a powerful model to study the genetic and neural mechanisms underlying the evolution of behaviors. *A. mexicanus* exists as two morphotypes: a sighted, surface-dwelling fish, and multiple cave-dwelling populations of blind fish. As cave populations adapted to their environment, they have repeatedly evolved various morphological, physiological, and behavioral changes, including reductions in eye size and pigmentation, increased adipose tissue storage and insulin resistance, and changes in multiple behaviors including reductions in schooling, response to stressors, and aggression (Burchards, Dölle, and Parzefall, 1985; Chin et al., 2018, 2020; Elipot, Hinaux, Callebert, and Rétaux, 2013; Jeffery, 2001; J. E. Kowalko, Rohner, Rompani, et al., 2013; Riddle et al., 2018; Rodriguez-Morales et al., 2022; Swaminathan, Xia, and Rohner, 2024; Wilkens, 1988; Xiong et al., 2022). Cave and surface morphs of *A. mexicanus* remain interfertile, making *A. mexicanus* a powerful model for studying the genetic underpinnings that contribute to the evolution of cave-evolved traits. Multiple genetic mapping studies on surface-cave hybrids have successfully identified the genetic architecture underlying many cave-evolved traits (J. E. Kowalko, Rohner, Linden, et al., 2013; J. E. Kowalko, Rohner, Rompani, et al., 2013; O’Quin, Yoshizawa, Doshi, and Jeffery, 2013; M. Protas, Conrad, Gross, Tabin, and Borowsky, 2007; M. E. Protas et al., 2006; M. Protas et al., 2008; Riddle et al., 2021; Warren et al., 2021; Yoshizawa et al., 2015; Yoshizawa, Yamamoto, O’Quin, and Jeffery, 2012). Moreover, through the employment of genetic tools such as CRISPR-Cas9, specific candidate genes identified through these mapping studies can now be manipulated to investigate function (Klaassen, Wang, Adamski, Rohner, and Kowalko, 2018; J. Kowalko, 2020; Sifuentes-Romero, Ferrufino, and Kowalko, 2023; Stahl et al., 2019). For example, these approaches have led to the functional assessment of genes associated with the evolution of eye development, including the *cystathionine B-synthase a (cbsa)* and *retinal homeobox 3 (rx3)* genes, and pigmentation, including the *oculocutaneous albinism 2 (oca2)* gene (Klaassen et al., 2018; Ma et al., 2020; Shennard et al., 2024). The genetic tractability of this species demonstrates that *A. mexicanus* is a powerful system in which to investigate the genetic underpinnings of evolved differences in behaviors.

*A. mexicanus* surface-dwelling morphs exhibit territorial aggression, which is crucial in establishing hierarchical dominance structures (Burchards et al., 1985; Elipot et al., 2013; Espinasa, Yamamoto, and Jeffery, 2005). In adult fish, aggression is composed of multiple behaviors, including striking, biting, following, circling, fin-spreading, and snake-swimming (Burchards et al., 1985; Rodriguez-Morales et al., 2022). Aggression has also been reported in younger *A. mexicanus* (3 months), however, studies in younger fish have focused solely on striking behavior (Elipot et al., 2013). While cavefish from some cave populations display aggression, fish from multiple cave populations show reductions in the aggression-associated behaviors striking and following relative to surface fish (Espinasa et al., 2022; Rodriguez-Morales et al., 2022). In addition, circling, which has been reported as an aggressive behavior in other fish species (Xu et al., 2021; Zabegalov et al., 2019), occurs more frequently in at least one cavefish population than in surface fish (Rodriguez-Morales et al., 2022). While aggression has been studied extensively in *A. mexicanus*, little is known about the genetic underpinnings of evolved differences in aggression-associated behaviors in *A. mexicanus*. A central challenge to applying genetic mapping and functional genetic approaches to behavioral studies is the need for accurate and consistent quantification of behaviors in large datasets. Previous aggression studies in *A. mexicanus* relied on manual annotations (Elipot et al., 2013; Rodriguez-Morales et al., 2022), but these methods are time intensive and require extensive training to standardize annotations between observers, making studies evaluating genetic underpinnings of aggressive behaviors challenging. To effectively investigate the genetic underpinnings of aggression, methods are needed to automate behavioral annotations in this species.

Recent advancements in machine learning tracking and behavioral analysis tools such as Simple Behavioral Analysis (SimBA) and Social LEAP Estimates Animal Pose (SLEAP) have provided researchers with robust methods for rapid, automated data analysis (Goodwin et al., 2024; Pereira et al., 2022). These machine-learning based tools significantly reduce manual annotation time, minimize bias between different annotators, and allow for the processing of large datasets to improve statistical power and reproducibility (Goodwin et al., 2024; Pereira et al., 2022). Here, we establish methods in *A. mexicanus* for automated tracking and behavioral analysis with the aim of elucidating the patterns of aggression in *A. mexicanus* juveniles and assessing genetic architecture underlying evolved differences in aggressive behaviors utilizing cave-surface hybrid fish. This pipeline lays the groundwork for identifying the loci and genes contributing to natural variation in aggression between surface and cavefish in this species.

## Methods

### Fish husbandry and specimens

*A. mexicanus* used in the study were derived from lab-bred populations of surface fish from Mexico and Pachón cavefish that had been maintained in the laboratory for multiple generations and raised following standardized protocols (Kozol et al., 2023), with slight modifications. Surface-cave F1 hybrids were generated by crossing a male surface fish with a female Pachón cavefish. The surface-Pachón hybrid F2 population (F2) was derived from a single pair of F1 siblings. Two F1 hybrids were crossed to generate a line of F2 hybrids. All adult, larval, and juvenile fish post 6 days post fertilization (dpf) were housed at 23±1°C on a 14:10 light:dark cycle. Larvae for aggression assays were raised in 250 mL of water at a density of 100 fish per bowl in incubators equipped with lights set to a 14:10 light dark cycle at 25°C. Fish were moved to 2L plastic tanks at 6 dpf. From 6 dpf to 29 dpf, fish were fed brine shrimp twice a day. At 14 dpf, fish were moved into 2L tanks on a recirculating Aquaneering system at a density of 15 fish per tank. Beginning at 30 dpf, fish were fed brine shrimp twice a day and gemma (GEMMA Micro 300 from Skretting) once a day. Fish were size sorted as needed to ensure relatively similar sizes between fish housed in the same tank. All experiments occurred at Lehigh University in accordance with Institutional Animal Care and Use Committees (IACUC) protocols.

### Resident/Intruder assays

Aggression was elicited using a resident-intruder assay (Elipot et al., 2013), with modifications. Once fish reached 1.4 cm, they were fed brine shrimp once a day for at least 1 week prior to the assay. Fish were food deprived for 3 days before assays were performed, as in (Burchards et al., 1985). All assayed fish were between 57-146 dpf and 1.4-2.5 cm in length. Assays were performed in water acclimated to an ambient room temperature of 25±1°C. Fish were placed in custom made 12x9x5 cm tanks filled with 350 mL of conditioned fish system water for at least 18 hours prior to assays between Zeitgeber (ZT) time ZT7-ZT9. Assay tanks were placed on a 14:10 custom made infrared and LED light boxes which kept a 14:10 light/dark cycle. Assays were performed in the light. The focal resident and non-focal intruder fish were intentionally paired so the resident was visibly larger than the intruder. After the acclimation period, the intruder fish was placed into the resident tank. All assays were conducted between ZT1-ZT7. Videos were recorded from above on a Basler ace acA1300-60gm or acA1280-60gm GigE Mono camera attached to 16mm C Series Lens, VIS-NIR (Edmund Optics) at 30 fps for 62 minutes. Videos were recorded using PylonViewer software (Basler).

### Multi-animal pose estimation

For automated multi-animal pose tracking, we utilized and trained the machine learning model Social LEAP Estimates Animal Poses (SLEAP) (Pereira et al., 2022) on an Ubuntu 20.04.6 LTS desktop equipped with 11^th^ Gen Intel® Core™ i7-11700F processor (2.50 GHz), 66.0 GB of RAM, and a NVIDIA GeForce RTX 3060 Ti Lite Hash Rate (LHR) graphics card to identify and track both resident and intruder fish during the behavioral assays. 9 nodes were assigned to key body-parts of each fish to compromise the project skeleton–including nose, head, left eye, right eye, upper body, center, lower body, tail, and fin (Supplemental Fig. 1). Body parts were manually labeled on a subset of frames from the training videos imported as grayscale. Training was initially performed using a multi-animal top-down pipeline, with the center of the fish as the anchor. The trained model was then run on 20 random frames, and model performance was assessed for accuracy by manually comparing computer generated pose-estimations to true body parts. Additional frames were manually annotated for the training dataset until performance was satisfactory based on manual inspection of the output of the training videos. The final model was trained on a total of 526 manually annotated frames across 1 surface-resident/Pachón-intruder, 2 surface-resident/surface-intruder, and 3 Pachón-resident/Pachón-intruder videos. All videos were run through the final trained model with the following parameters: Tracking_pipeline=multi-animal top-down, max_instances=2, batch_size=4, tracking.tracker=flow, tracking.max_tracking=2, tracking.similarity=instance, and tracking.match=greedy. The tracks were exported as (.h5) files for analysis.

### Manual behavior annotations

Behaviors for the training and test datasets were manually annotated and cross-referenced by two trained annotators using Behavioral Observation Research Interactive Software (BORIS) (Friard and Gamba, 2016). Training videos were split into 10-minute segments and manually annotated for striking, circling, and following behaviors as state events. A total of 34 10-minute videos were manually annotated for the striking training dataset, consisting of 8 surface-resident/surface-intruder, 9 surface-resident/Pachón-intruder, and 17 Pachón-resident/ Pachón-intruder assay videos. For the following and circling training dataset, a total of 30 10-minute videos were manually annotated, consisting of 6 surface-resident/surface-intruder, 9 surface-resident/Pachón-intruder, and 15 Pachón-resident/ Pachón-intruder assay videos. 12 10-minute videos were manually annotated for the test dataset, consisting of 6 surface-resident/surface-intruder and 6 Pachón-resident/ Pachón-intruder assay videos. The training and test videos chosen were included in the parental population dataset. Final annotations for each video were exported as a (.csv) file for analysis.

### Automated quantification of behavior

Classifiers were built using Simple Behavioral Analysis (SimBA) algorithms for automated quantification of behaviors (Goodwin et al., 2024). SimBA project configurations were created and executed on a Windows 11 Enterprise (64-bit) computer equipped with a 13th Gen Intel® Core™ i7-13700 processor (2.10 GHz), 16.0 GB of RAM, and an Intel® UHD Graphics 770 GPU. Since following behaviors may occur simultaneously with striking behavior, a separate project configuration file was created for following, while striking and circling models exist in a single project. Project configuration files specified the number of behavioral classifiers per project, and a custom multi-animal body-part configuration was created to mirror SLEAP pose-estimation with 9 body parts per fish. Videos (.mp4), SLEAP tracking files (.h5), and BORIS training files (.csv) were imported for each training video. Video parameters were indicated and pixels per millimeter was defined by using the length of the tank (120mm). Outlier correction was skipped, and features were extracted for each video. We developed behavior classifiers using a random forest model. The model was implemented with parameters including 2,000 random forest estimators, a minimum sample leaf node of 1, RF_criterion set to "gini," RF_max_features set to "sqrt," and a test size of 20%. Minimum bout durations were determined based on manual annotations, with thresholds set at 67 ms for striking, 200 ms for following, and 300 ms for circling.

A precision-recall performance analysis curve was generated for each classifier during training to assess model performance (Supplemental Fig. 2). The harmonic mean between precision and recall was reported as the F1 curve, where F1_max_ corresponds to the optimal discrimination threshold value, or the probability cutoff in determining if the behavior is present. The discrimination value was manually adjusted to yield optimal performance for striking=0.375, following=0.275, and circling=0.174 based on manual inspection of videos at a range of discrimination thresholds. Classifier models were run on the unseen testing dataset and compared to the manual annotations to assess performance. Performance was considered satisfactory by a trained experimenter who manually inspected SimBA’s output in comparison to the manual annotations for all training and test videos. All videos were run through the pipeline, predicting bout number (striking and circling) and total bout duration (s) (following). To further ensure SimBA correctly identified behaviors, and to determine if behaviors were performed by F2 fish or cavefish in the F2 behavioral analysis, 10 SimBA annotated behaviors across 10 F2 videos were manually inspected. For striking, 79 of 100 behaviors were accurate, with 79 of 79 strikes being performed by the resident F2. Following was occurring for 98 of 100 behavioral timepoints indicated by SimBA, with 98 of 98 being performed by the resident F2. Due to a limited total number of circling events through the videos, only 10 behavioral timepoints were analyzed, with 8 of 10 behaviors being accurately annotated by SimBA.

### Quantification of eye size and standard length

Post assay, F2 resident fish were euthanized and imaged on a Canon Rebel DSLR camera for standard length and eye size. Standard length, measured from the nose to the end of the body, and eye perimeter quantifications were performed in Fiji (Schindelin et al., 2012). Fish with partially occluded eyes or missing images (*n=*3) were excluded from eye-perimeter related statistical analyses.

### Statistical analysis

All statistical analyses were performed and graphs were generated using Graphpad Prism version 10.3.0, Graphpad Software, graphpad.com. Relative eye size was calculated by dividing the eye perimeter (mm) by standard length (mm), as eye size co-varies with standard length at this stage (Supplemental figure 3D,E). Spearman’s correlation coefficients were calculated to compare manual and automated quantification of behavior across all behavioral classifiers for the training and test datasets, except in Pachón following comparisons, in which Pearson’s correlation coefficient was calculated as the data was normally distributed. Mann-Whitney tests were used to compare behavior in all the parental surface and Pachón fish vs. control fish of the same population. Kruskal-Wallis with Dunn’s post-hoc multiple comparisons tests were used to assess behavioral differences between surface, Pachón, and F2 hybrid fish versus Pachón intruders. Spearman’s correlation coefficients were calculated to compare each of the behaviors to each other, as well as to the standard length and eye size. A multiple linear regression model was used to analyze the relationship between striking and following behaviors while statistically controlling for standard length.

## Results

### Development of a pipeline for automated quantification of multiple-stress associated behaviors

While previous research has demonstrated distinct differences in aggressive behaviors between surface-dwelling and cave-dwelling morphs (Burchards et al., 1985; Elipot et al., 2013; Espinasa et al., 2022, 2005; Rodriguez-Morales et al., 2022), these studies have relied on manual annotations of aggressive behaviors, which are time-intensive and require significant training to maintain reproducible annotations. To address this, we established a machine learning-based pipeline to enable automated tracking and quantification of these behaviors in juvenile fish (Fig. 1A). To accurately track behavior, we modified the standard resident-intruder assay that has been used to induce aggression in *A. mexicanus* (Elipot et al., 2013). To optimize tracking, fish were filmed from above on a background that was backlit with white light and infrared light. Assays were performed on juvenile (57-146 dpf) fish to minimize space needed for assays and to reduce time needed to raise and house fish on the system.

**Figure 1.**
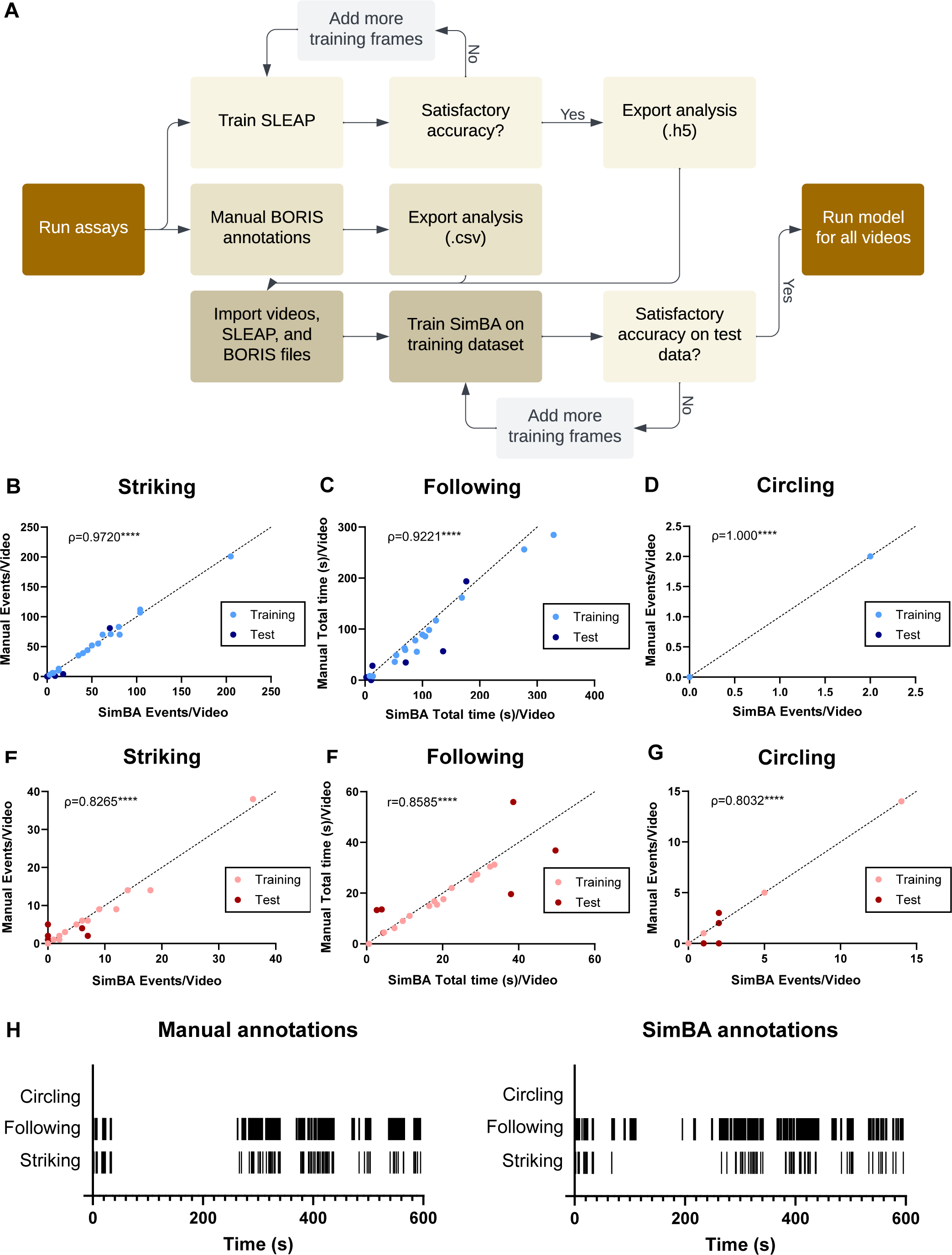
Automated tracking pipeline and performance analysis. (A) Automated behavioral analysis pipeline. Resident-Intruder assays were performed and recorded. A trained and validated SLEAP model was used to perform animal pose-estimation for each video and exported as an .h5 file. Manual annotations for 3 aggressive behaviors were completed in BORIS and used for training the behavioral classifiers in SimBA. The trained models were assessed against training (striking: *n*=34, following and circling *n=*30) and novel (*n*=12) videos to demonstrate satisfactory performance was achieved. All videos were run through SLEAP and SimBA for automated tracking and behavioral analysis for each of the classifiers. (B-G) Classifier performance analysis. Spearman’s correlation (A-D,F) and Pearson’s correlation (E) between manual and SimBA automated annotations on the training and novel testing dataset in surface fish for (B) striking (*rho*=0.9720, *p*<0.0001), (C) following (*rho*=0.9221, *p*<0.0001), and (D) circling (*rho*=1.000, *p*<0.0001), and Pachón cavefish for (E) striking (*rho*=0.8265, *p*<0.0001), (F) following (*r*=0.9247, *p*<0.0001), and (G) circling (*rho*=0.8032, *p*<0.0001). (H) Ethogram depicting manually annotated events for all 3 behaviors on a novel surface fish video compared to SimBA’s annotations. All testing and training videos were 10 minutes in length.

To establish our automated pipeline, training and test videos were obtained by performing surface-resident/surface-intruder, surface-resident/ Pachón-intruder, and Pachón-resident/Pachón-intruder assays. Pose-estimation was performed using SLEAP software, with each fish tracked across nine defined body nodes: nose, head, left eye, right eye, upper body, center, lower body, tail, and fin (Supplemental Fig. 1). Model efficiency was assessed by a trained experimenter through manual inspection of SLEAP generated pose-estimations. By comparing these to the true body parts of each fish until SLEAP accurately labeled each body part, including in obscured frames, we established that our model performed accurate labeling of fish with 526 frames manually annotated.

To develop automated methods for behavioral annotations, we performed manual annotations of behaviors in sets of videos to train and test the accuracy of our pipeline. Each video recording in the training and test datasets was imported into BORIS for manual annotation of three aggression-associated behaviors previously identified in adult surface fish: following, striking, and circling (Rodriguez-Morales et al., 2022). The videos, manually annotated behavioral data from the training dataset, and SLEAP tracking files were imported into the machine-learned program, SimBA, to train behavior classifiers for striking, following, and circling. After identifying the optimal discrimination threshold values, we determined how accurately the models performed. The trained striking classifier was run on the training dataset, consisting of 8 surface-resident/surface-intruder, 9 surface-resident/Pachón-intruder, and 17 Pachón-resident/Pachón-intruder 10-minute videos. Following and circling trained classifiers were run on the training dataset, consisting of 6 surface-resident/surface-intruder, 9 surface-resident/Pachón-intruder, and 15 Pachón-resident/Pachón-intruder 10-minute videos. All the classifiers were then run on the unseen test datasets, consisting of 6 surface-resident/surface-intruder and 6 Pachón-resident/Pachón-intruder 10-minute videos. Their outputs were compared to manual annotations for both surface and Pachón populations to evaluate model performance and confirm the absence of population bias. We found strong correlations between manual annotations and SimBA derived annotations for both cave and surface fish for striking (Fig. 1B, E), following (Fig. 1C, F) and circling (Fig. 1D, G), suggesting that the SLEAP/SimBA pipeline for automated behavioral analysis is efficient and accurate at scoring all three aggressive behaviors across *A. mexicanus* populations. Moreover, ethograms of behaviors show that the patterns observed in manually annotated videos and SimBA annotated videos are largely similar, further supporting the use of the automated pipeline for annotating behavior (Fig. 1H).

### Multiple aggression associated behaviors are altered in juvenile Pachón cavefish

To determine if there are quantitative, morph-specific differences in aggressive behaviors in juvenile fish, resident-intruder assays were conducted using either surface-dwelling (*n*=10) or Pachón cave-dwelling (*n*=10) pairs, in which both the resident and intruder originated from the same population. Juvenile surface fish exhibited a significantly higher number of striking bouts and more time following compared to Pachón counterparts (Fig. 2A,B). In contrast, Pachón cavefish display a significantly greater number of circling events (Fig. 2C), suggesting that previously reported distinctions in aggressive phenotypes between these morphs in adult animals (Rodriguez-Morales et al., 2022) are also present in juvenile animals. Previous work found that attacks in surface fish occur more frequently at later timepoints in the resident-intruder assay, whereas attacks in Pachón cavefish occur earlier in the assay (Elipot et al., 2013). To determine if the behaviors quantified here occur at different timepoints between morphs, we binned behaviors into five-minute increments, and plotted when these behaviors occur across the time of the assay. We found qualitative differences in when striking occurs during the assay in surface and cavefish with surface fish striking more in the second half of the video and cavefish striking more in the first half of the video (Fig. 2B), consistent with the patterns of attack frequency previously reported in these populations (Elipot et al., 2013).

**Figure 2.**
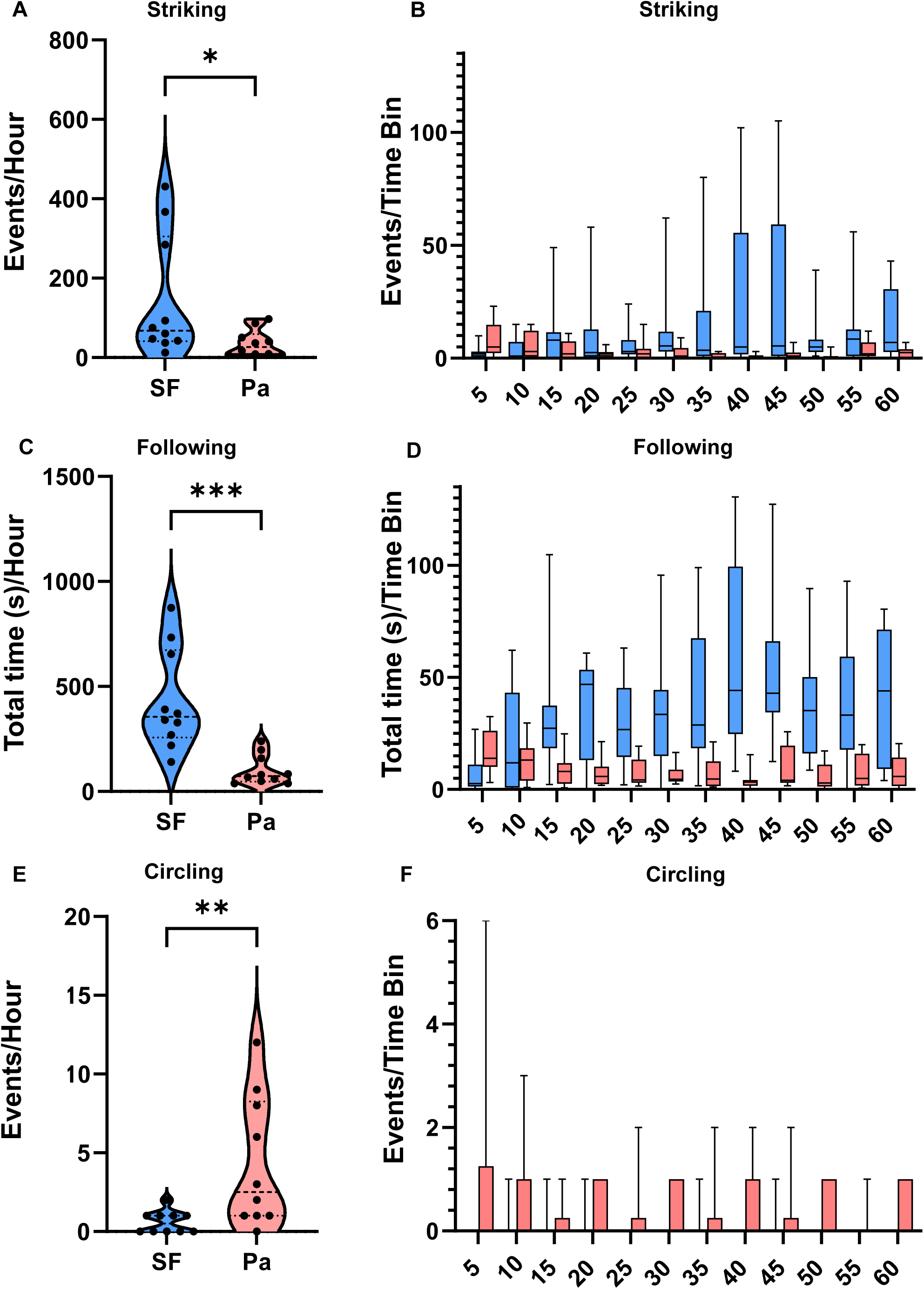
Quantification and temporal analysis of aggressive behaviors in parental populations. Comparison of aggression-associated behaviors in surface-resident/surface-intruder (*n=*10) and Pachón resident/Pachón intruder assays (*n=*10). Mann-Whitney U-tests were performed for each behavior to compare the two groups. Significant differences were observed for (A) striking (*U*=21, *p*=0.0281), (C) total time spent following (*U=*4, *p*=0.0001), and (E) circling bouts (*U=*17, *p*=0.0096). (Pa = Pachón, SF = surface fish). These behaviors were quantified across 5-minute time bin intervals for (B) striking, (D) following (s), and (F) circling bouts per hour.

Similarly, surface fish performed following more often in the second half of the video compared to the first half (Fig. 2D). In contrast, following in cavefish and circling in both populations did not show differences in patterns across the time of the assay (Fig. 2F).

### Analysis of surface-Pachón hybrids reveals co-variance of aggression associated traits

Surface fish and Pachón cavefish are interfertile, making *A. mexicanus* a powerful model for investigating the genetic basis of complex traits. Assessing multiple traits in F2 hybrids allows for analysis of trait co-variation, providing insight into whether traits evolved through shared genetic loci due to the same genes or close linked genes, or independently. We applied the automated pipeline to quantify aggression-associated behaviors in F2 hybrids to elucidate the genetic architecture underlying these traits, as well as the relationships between these traits, and their relationships with morphological traits. To determine individual fish behavior, we assayed the resident focal fish (surface, Pachón, or surface-Pachón F2 hybrid fish) with a smaller Pachón intruder. Surface fish (*n*=14) performed more striking bouts and spent more time following compared to Pachón fish (*n*=14), with following reaching statistical significance (Fig. 3A,B). Conversely, Pachón fish exhibited an increased number of circling bouts compared to surface fish, although this difference did not reach statistical significance (Fig. 3C). This suggests that surface fish maintain aggression against a Pachón intruder, while Pachón fish do not demonstrate significant following or striking in the assay but will elicit circling, which requires two participating fish. Together, these results suggest that using this method of assaying F2s against a Pachon intruder fish will allow for accurate quantification of F2 aggressive profiles.

**Figure 3.**
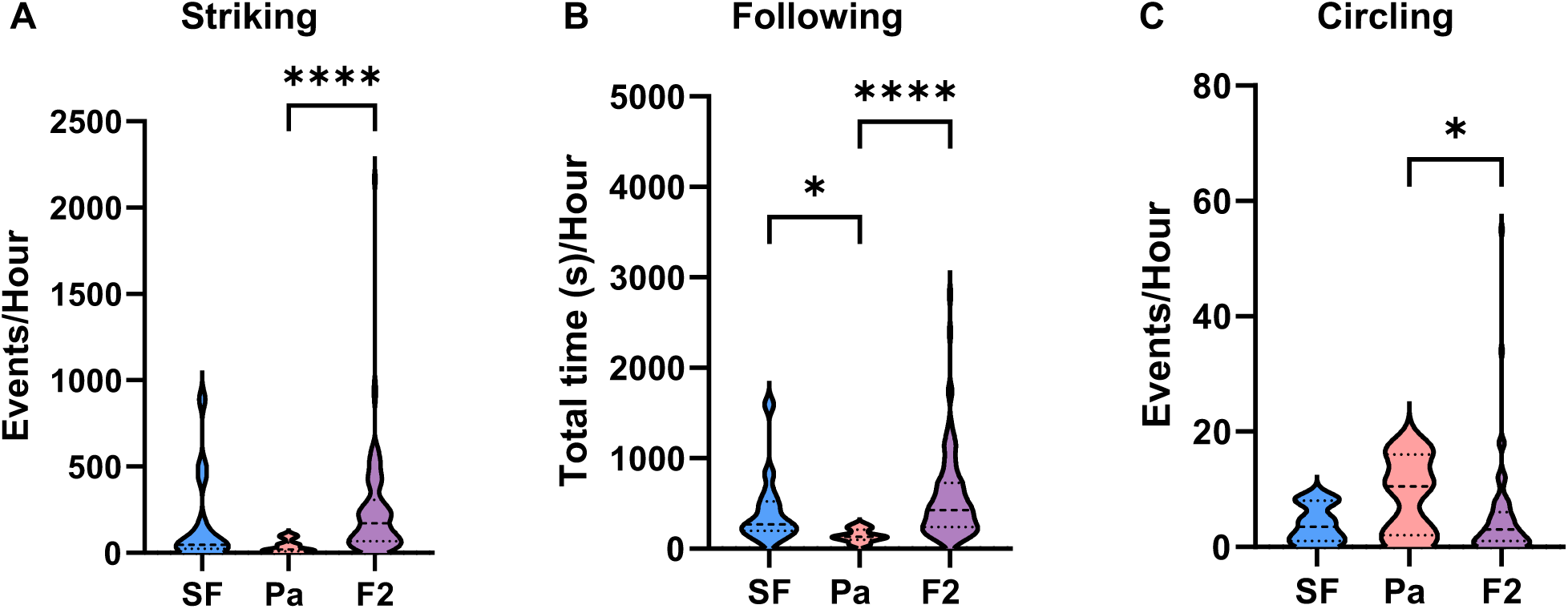
Quantification of aggression in parentals and F2 hybrids. Comparison of (A) number of strikes, (B) total time spent following (s), and (C) circling bouts among surface (*n=*14), Pachón (*n=*14), and F2 hybrids (*n*=97) assayed against a Pachón intruder. Kruskal-Wallis tests followed by Dunn’s post-hoc multiple comparisons were performed. Statistical comparisons for each behavior are as follows: (A) Striking (*H(2)=*24.72): surface vs. Pachón (*p*=0.2036), surface vs. F2 (*p*=0.0708), Pachón vs. F2 (*p*<0.0001), (B) following (*H(2)=*24.50): surface vs. Pachón (*p*=0.0210), surface vs. F2 (*p*=0.5485), Pachón vs. F2 (*p<*0.0001), and (C) circling (*H(2)*=5.848): surface vs. Pachón (*p*=0.3854), surface vs. F2 (*p* >0.9999), Pachón vs. F2 (*p*<0.0471). (Pa = Pachón, SF = surface fish, F2 = surface-Pachón F2).

Indeed, manual inspection of a subset of F2 behaviors confirmed that the focal F2 fish was the primary instigator of aggressive behaviors (see Methods), suggesting that this method of quantifying F2 social behaviors is robust. F2 hybrids (*n*=97) displayed a broad distribution of phenotypes ranging from surface-like to cave-like across all three behaviors (Fig. 3A-C). This suggests that aggression is genetically encoded, and F2 hybrid animals can be used to investigate the genetic underpinnings of the evolution of aggressive behaviors in this species.

We next sought to determine whether any morphological traits co-vary with aggression-associated traits in this population. Standard length was significantly positively correlated with both striking and following in F2 fish (Fig. 4A,B). This relationship was not due to larger fish being older, as striking and following were both not correlated with fish age (Supplemental Fig. 3G,H). Circling, in contrast, was not significantly correlated with standard length or fish age (Fig. 4C, Supplemental Fig. 3G-I). Next, we assessed whether eye size correlates with any aggressive behaviors, as vision is required for another social behavior, schooling (J. E. Kowalko, Rohner, Rompani, et al., 2013; Patch et al., 2022). Eye size and following behavior were moderately correlated (Fig. 4E). Striking and circling, however, were not correlated with eye size (Fig. 4D, F), suggesting that some aggression-associated behaviors evolved independently from eye size, and consistent with previous results suggesting that aggression is not dependent on having visual cues (Rodriguez-Morales et al., 2022).

**Figure 4.**
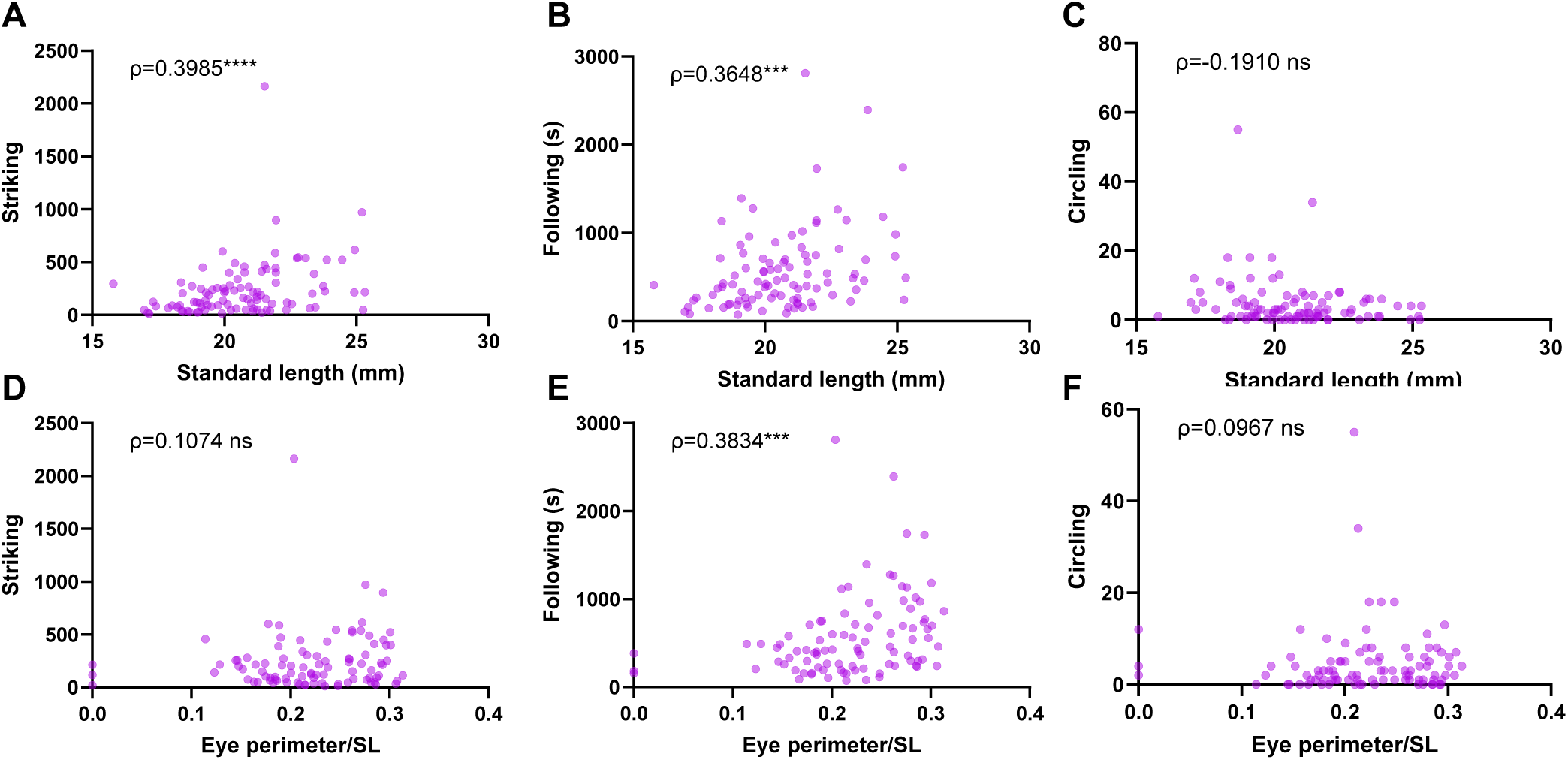
Correlations between morphology and aggression behaviors in F2 hybrids. Spearman correlation in F2 hybrids (*n=*96) between standard length (mm) and (A) striking (*rho*=0.3985, *p*<0.0001), (B) following (s) (*rho*=0.3648*, p*=0.0002), and (C) circling bouts (*rho=*-0.1910*, p*=0.0609) per hour. Spearman correlation in F2 hybrids (*n=*94) between right eye perimeter (mm) corrected for standard length and (D) striking (*rho*=0.1074, *p*=0.3030), (E) following (s) (*rho*=0.3834, *p=*0.0001), and (F) circling bouts (*rho*=0.0967, *p*=0.3538) per hour.

To investigate whether these aggression-associated behaviors evolved independently, we assessed correlations among these behaviors within the F2 hybrid population. Striking and following behaviors are strongly positively correlated (Fig. 5A), suggesting that they may evolve through a shared genetic basis. As both striking and following behaviors were positively correlated with standard length, a multiple linear regressions model was employed to confirm the relationship between striking and following behaviors occurred independently of standard length. The model showed the total time spent of following was an accurate predictor of striking bouts (*t*=12.02, *p*<0.0001), while standard length was not (*t*=0.7795, *p*=0.4378). Conversely, striking and circling show a significant, moderate negative correlation, suggesting that at least some shared genetic factors may contribute to the evolution of these traits (Fig. 5B). No significant correlation was observed between following and circling behaviors (Fig. 5C), indicating these traits may evolve through independent genetic factors. Together, these results offer insight into the genetic mechanisms by which aggression-associated behaviors evolved in *A. mexicanus*.

**Figure 5.**
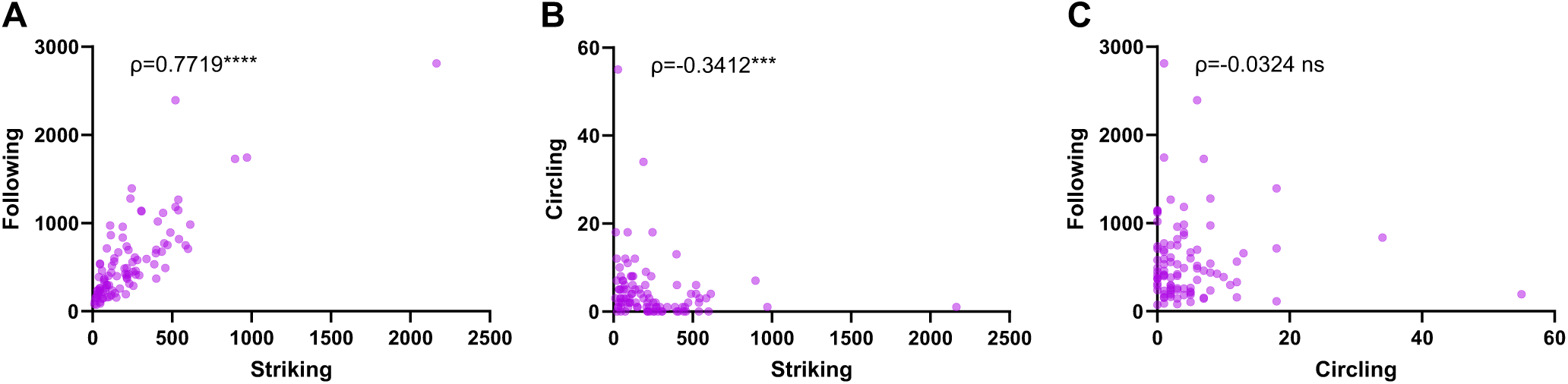
Correlations between aggressive behaviors in F2 hybrids. Spearman correlations between behaviors per hour in F2 hybrids (*n=*97) between (A) number of strikes and total time spent following (s) (*rho*=0.7719, *p*<0.0001), (B) number of strikes and total circling bouts (*rho*=0.3412, *p*=0.0006), and (C) total circling bouts and total time spent following (s) (*rho*=-0.0324, *p=*0.7531).

## Discussion

### Machine-learning algorithms for phenotyping aggression-associated behaviors in Astyanax mexicanus

Behavioral neuroscience and genetics are rapidly growing fields that require succinct, accurate methods for behavioral quantification. However, manual quantification of behavior presents many challenges, as it is time-consuming, requires extensive training, and can introduce bias between different observers. Through the implementation of machine-learning based algorithms for automated tracking and behavioral quantification, these challenges can be addressed (Anderson and Perona, 2014; Datta, Anderson, Branson, Perona, and Leifer, 2019; Egnor and Branson, 2016; Goodwin et al., 2024; Pereira et al., 2022). This study establishes and validates a machine learning-based pipeline for the automated quantification of aggression-associated behaviors in *A. mexicanus*. By replacing manual behavioral annotations with an automated machine-learning approach, we demonstrate the feasibility of rapid, robust quantification of complex behaviors.

While machine learning based methods have recently been established to assess behaviors in individually assayed fish in *A. mexicanus* (Padmanaban et al., 2025), this work provides a significant advancement by providing methods to assess social behaviors with multiple individuals in this species. Automated tracking and analysis of social behaviors presents various challenges related to tracking multiple animals at once due to factors such as identity switching or obscurity due to overlapping animals. Although similar methods have been used to quantify social behaviors in other species like *Rattus norvegicus* (Lapp, Salazar, and Champagne, 2023), implementing this pipeline in *A. mexicanus* significantly advances the field’s capacity to perform high-throughput phenotyping in an established model for evolutionary genetics (Casane and Rétaux, 2016; J. Kowalko, 2020). This provides the foundation to considerably accelerate the completion of larger genetic mapping studies for aggression, that were limited by the labor-intensive nature of manual annotations. Through these genetic mapping studies, we can identify aggression-associated candidate genes and utilize gene editing methods that have been optimized in the species, such as CRISPR-Cas9(Klaassen et al., 2018; Sifuentes-Romero et al., 2023; Stahl et al., 2019), to manipulate their function and assess their impact on aggression. Additionally, methods established in this study for rapid, controlled quantification of aggressive behavior can be adapted to assess aggression at other developmental timepoints, as well as to assess other social behaviors that vary in *A. mexicanus* such as schooling and food competition (Choy et al., 2025; Espinasa, Bibliowicz, Jeffery, and Rétaux, 2014; J. E. Kowalko, Rohner, Rompani, et al., 2013).

### Juvenile A. mexicanus display similar behavioral changes to those observed in mature adults

Our findings confirm and build on previous observations that surface fish juveniles display significantly more aggressive striking behaviors than cavefish (Elipot et al., 2013). While we replicated this finding, we also expanded on this to investigate behavioral differences in other aggression-associated behaviors previously identified in adults, following and circling (Rodriguez-Morales et al., 2022). Here, we showed juvenile surface fish perform significantly more following compared to Pachón, while Pachón showed an increased tendency for circling. These results are similar to what was previously reported in adult fish (Rodriguez-Morales et al., 2022), demonstrating that younger individuals can be used for these assays. Earlier phenotyping is beneficial for larger genetic mapping studies to preserve resources, labor, and space spent raising and housing individuals. While we did not find that fish age played a large role in aggressive behaviors in our dataset, we did find a positive correlation between aggressive associated behaviors and standard length, suggesting that future studies should ensure fish are within a specific size range for these assays.

### Co-variance in F2 hybrids suggests a role for shared genetic underpinnings of different aggression-associated behaviors and morphological traits

In the F2 hybrid population, we observed a broad distribution of behavioral phenotypes ranging from surface-like to cave-like, suggesting a complex genetic architecture. However, significant correlations between specific behaviors–specifically a strong positive correlation between striking and following, and the negative relationship between striking and circling– suggest potential shared genetic underpinnings of these traits. Interestingly, we had previously identified circling as a behavior that is increased in only one of three independently evolved cave populations relative to surface fish (Rodriguez-Morales et al., 2022). While the increased tendency to circle in Pachón cavefish alone suggests that circling may be unrelated to the evolved changes in other aggression-associated behaviors, the observation that striking and circling co-vary in F2 hybrid fish suggests at least some relationship between these traits in this population.

The role of vision in the evolution of social behaviors in *A. mexicanus* has been studied extensively. Vision is not required for aggressive behaviors such as striking and circling. Surface fish populations maintain aggression in the dark, and some blind cavefish populations such as those from the Molino cave still perform strikes (Espinasa et al., 2022, 2005; Rodriguez-Morales et al., 2022). For other social behaviors such as schooling, *A. mexicanus* populations rely on vision (J. E. Kowalko, Rohner, Rompani, et al., 2013; Patch et al., 2022). Interestingly, eye size was significantly associated with following behavior but not with striking or circling in F2 hybrids. Given the known link between eye size and schooling behavior in *A. mexicanus*, this finding suggests that visual input may specifically influence social tracking or coordination–which is inherently integral to following behavior–rather than aggression alone. This distinction points to potentially distinct genetic pathways underlying aggression versus schooling, despite some overlapping sensory components.

## Conclusion

In summary, this study provides a powerful, unbiased alternative to the time-consuming nature of manual behavioral annotation for social behaviors in a model with evolved differences in multiple social behaviors, and demonstrates its efficiency in investigating the genetic basis of complex social traits. This pipeline sets the stage for future large-scale genetic mapping efforts and functional studies aimed at understanding the evolution of and mechanisms underlying aggression and related behaviors.

## Acknowledgements

This work was funded by National Institutes of Health grant R35GM138345.

## Notes

### Competing Interest Statement

The authors have declared no competing interest.

## References

Alonso, F., Cánepa, M., Moreira, R. G., and Pandolfi, M. (2011). Social and reproductive physiology and behavior of the Neotropical cichlid fish Cichlasoma dimerus under laboratory conditions. Neotropical Ichthyology, 9(3), 559–570.

Anderson, D. J., and Perona, P. (2014). Toward a science of computational ethology. Neuron, 84(1), 18–31.

Archer, J. (1988). The Behavioural Biology of Aggression. Cambridge, England: Cambridge University Press.

Burchards, H., Dölle, A., and Parzefall, J. (1985). Aggressive behaviour of an epigean population of Astyanax mexicanus (Characidae, Pisces) and some observations of three subterranean populations. Behavioural Processes, 11(3), 225–235.

Butler, K. (1973). Predatory behavior in laboratory mice: strain and sex comparisons. Journal of Comparative and Physiological Psychology, 85(2), 243–249.

Casane, D., and Rétaux, S. (2016). Evolutionary genetics of the cavefish Astyanax mexicanus. Advances in Genetics, 95, 117–159.

Chin, J. S. R., Gassant, C. E., Amaral, P. M., Lloyd, E., Stahl, B. A., Jaggard, J. B., … Duboue, E. R. (2018). Convergence on reduced stress behavior in the Mexican blind cavefish. Developmental Biology, 441(2), 319–327.

Chin, J. S. R., Loomis, C. L., Albert, L. T., Medina-Trenche, S., Kowalko, J., Keene, A. C., and Duboué, E. R. (2020). Analysis of stress responses in Astyanax larvae reveals heterogeneity among different populations. *Journal of Experimental Zoology. Part B*, Molecular and Developmental Evolution, 334(7–8), 486–496.

Choy, S., Thakur, S., Polyakov, E., Abdelaziz, J., Lloyd, E., Enriquez, M., … Kowalko, J. E. (2025). Mutations in the albinism gene oca2 alter vision-dependent prey capture behavior in the Mexican tetra. The Journal of Experimental Biology, 228(7), jeb249881.

Datta, S. R., Anderson, D. J., Branson, K., Perona, P., and Leifer, A. (2019). Computational neuroethology: A call to action. Neuron, 104(1), 11–24.

Egnor, S. E. R., and Branson, K. (2016). Computational analysis of behavior. Annual Review of Neuroscience, 39, 217–236.

Elipot, Y., Hinaux, H., Callebert, J., and Rétaux, S. (2013). Evolutionary shift from fighting to foraging in blind cavefish through changes in the serotonin network. Current Biology: CB, 23(1), 1–10.

Espinasa, L., Bibliowicz, J., Jeffery, W. R., and Rétaux, S. (2014). Enhanced prey capture skills in Astyanax cavefish larvae are independent from eye loss. EvoDevo, 5, 35.

Espinasa, L., Collins, E., Ornelas García, C. P., Rétaux, S., Rohner, N., and Rutkowski, J. (2022). Divergent evolutionary pathways for aggression and territoriality in Astyanax cavefish. Subterranean Biology, 43, 169–183.

Espinasa, L., Yamamoto, Y., and Jeffery, W. R. (2005). Non-optical releasers for aggressive behavior in blind and blinded Astyanax (Teleostei, Characidae). Behavioural Processes, 70(2), 144–148.

Friard, O., and Gamba, M. (2016). BORIS: a free, versatile open-source event-logging software for video/audio coding and live observations. Methods in Ecology and Evolution, 7(11), 1325–1330.

Goodwin, N. L., Choong, J. J., Hwang, S., Pitts, K., Bloom, L., Islam, A., … Golden, S. A. (2024). Simple Behavioral Analysis (SimBA) as a platform for explainable machine learning in behavioral neuroscience. Nature Neuroscience, 27(7), 1411– 1424.

Huntingford, F., and Turner, A. (1987). Animal Conflict.

Jeffery, W. R. (2001). Cavefish as a model system in evolutionary developmental biology. Developmental Biology, 231(1), 1–12.

Klaassen, H., Wang, Y., Adamski, K., Rohner, N., and Kowalko, J. E. (2018). CRISPR mutagenesis confirms the role of oca2 in melanin pigmentation in Astyanax mexicanus. Developmental Biology, 441(2), 313–318.

Kowalko, J. (2020). Utilizing the blind cavefish Astyanax mexicanus to understand the genetic basis of behavioral evolution. The Journal of Experimental Biology, 223(Pt Suppl 1). 10.1242/jeb.208835

Kowalko, J. E., Rohner, N., Linden, T. A., Rompani, S. B., Warren, W. C., Borowsky, R., … Yoshizawa, M. (2013). Convergence in feeding posture occurs through different genetic loci in independently evolved cave populations of *Astyanax mexicanus*. Proceedings of the National Academy of Sciences of the United States of America, 110(42), 16933–16938.

Kowalko, J. E., Rohner, N., Rompani, S. B., Peterson, B. K., Linden, T. A., Yoshizawa, M., … Tabin, C. J. (2013). Loss of schooling behavior in cavefish through sight-dependent and sight-independent mechanisms. Current Biology: CB, 23(19), 1874–1883.

Kozol, R. A., Yuiska, A., Han, J. H., Tolentino, B., Lopatto, A., Lewis, P., … Duboué, E. R. (2023). Novel husbandry practices result in rapid rates of growth and sexual maturation without impacting adult behavior in the blind Mexican cavefish. Zebrafish, 20(2), 86–94.

Lapp, H. E., Salazar, M. G., and Champagne, F. A. (2023). Automated maternal behavior during early life in rodents (AMBER) pipeline. Scientific Reports, 13(1), 18277.

Ma, L., Gore, A. V., Castranova, D., Shi, J., Ng, M., Tomins, K. A., … Jeffery, W. R. (2020). A hypomorphic cystathionine ß-synthase gene contributes to cavefish eye loss by disrupting optic vasculature. Nature Communications, 11(1), 2772.

Moyer, K. E. (1968). Kinds of Aggression and their Physiological Basis. Original Articles [Communications in Behavioral Biology. Part A], 2(2), 65–87.

O’Quin, K. E., Yoshizawa, M., Doshi, P., and Jeffery, W. R. (2013). Quantitative genetic analysis of retinal degeneration in the blind cavefish Astyanax mexicanus. PloS One, 8(2), e57281.

Padmanaban, N., Ambosie, R., Choy, S., Marcus, S., Nilsson, S. R. O., Keene, A. C., … Duboué, E. R. (2025). Automated behavioral profiling using neural networks reveals differences in stress-like behavior between cave and surface-dwelling Astyanax mexicanus (p. 2025.01.30.635725). 10.1101/2025.01.30.635725

Patch, A., Paz, A., Holt, K. J., Duboué, E. R., Keene, A. C., Kowalko, J. E., and Fily, Y. (2022). Kinematic analysis of social interactions deconstructs the evolved loss of schooling behavior in cavefish. PloS One, 17(4), e0265894.

Pereira, T. D., Tabris, N., Matsliah, A., Turner, D. M., Li, J., Ravindranath, S., … Murthy, M. (2022). SLEAP: A deep learning system for multi-animal pose tracking. Nature Methods, 19(4), 486–495.

Protas, M., Conrad, M., Gross, J. B., Tabin, C., and Borowsky, R. (2007). Regressive evolution in the Mexican cave tetra, Astyanax mexicanus. Current Biology: CB, 17(5), 452–454.

Protas, M. E., Hersey, C., Kochanek, D., Zhou, Y., Wilkens, H., Jeffery, W. R., … Tabin, C. J. (2006). Genetic analysis of cavefish reveals molecular convergence in the evolution of albinism. Nature Genetics, 38(1), 107–111.

Protas, M., Tabansky, I., Conrad, M., Gross, J. B., Vidal, O., Tabin, C. J., and Borowsky, R. (2008). Multi-trait evolution in a cave fish, Astyanax mexicanus. Evolution & Development, 10(2), 196–209.

Riddle, M. R., Aspiras, A. C., Gaudenz, K., Peuß, R., Sung, J. Y., Martineau, B., … Rohner, N. (2018). Insulin resistance in cavefish as an adaptation to a nutrient-limited environment. Nature, 555(7698), 647–651.

Riddle, M. R., Aspiras, A., Damen, F., McGaugh, S., Tabin, J. A., and Tabin, C. J. (2021). Genetic mapping of metabolic traits in the blind Mexican cavefish reveals sex-dependent quantitative trait loci associated with cave adaptation. BMC Ecology and Evolution, 21(1), 94.

Rodriguez-Morales, R., Gonzalez-Lerma, P., Yuiska, A., Han, J. H., Guerra, Y., Crisostomo, L., … Kowalko, J. E. (2022). Convergence on reduced aggression through shared behavioral traits in multiple populations of Astyanax mexicanus. BMC Ecology and Evolution, 22(1), 116.

Ruby, D. E. (1978). Seasonal changes in the territorial behavior of the iguanid lizard Sceloporus jarrovi. Copeia, 1978(3), 430.

Sands, J., and Creel, S. (2004). Social dominance, aggression and faecal glucocorticoid levels in a wild population of wolves, Canis lupus. Animal Behaviour, 67(3), 387– 396.

Schenkel, R. (1967). Submission: Its features and function in the wolf and dog. Integrative and Comparative Biology, 7(2), 319–329.

Schindelin, J., Arganda-Carreras, I., Frise, E., Kaynig, V., Longair, M., Pietzsch, T., … Cardona, A. (2012). Fiji: an open-source platform for biological-image analysis. Nature Methods, 9(7), 676–682.

Shennard, D., Sifuentes-Romero, I., Ambosie, R., Abdelaziz, J., Duboue, E. R., and Kowalko, J. E. (2024). The rx3 gene contributes to the evolution of eye loss in the cavefish Astyanax mexicanus (p. 2024.12.11.628050). 10.1101/2024.12.11.628050

Sifuentes-Romero, I., Ferrufino, E., and Kowalko, J. E. (2023). Application of CRISPR-Cas9 for Functional Analysis in A. mexicanus. In Neuromethods. Neuromethods (pp. 193–220). New York, NY: Springer US.

Stahl, B. A., Jaggard, J. B., Chin, J. S. R., Kowalko, J. E., Keene, A. C., and Duboué, E. R. (2019). Manipulation of Gene Function in Mexican Cavefish. Journal of Visualized Experiments: JoVE, (146). 10.3791/59093

Swaminathan, A., Xia, F., and Rohner, N. (2024). From darkness to discovery: evolutionary, adaptive, and translational genetic insights from cavefish. Trends in Genetics: TIG, 40(1), 24–38.

Warren, W. C., Boggs, T. E., Borowsky, R., Carlson, B. M., Ferrufino, E., Gross, J. B., … Rohner, N. (2021). A chromosome-level genome of Astyanax mexicanus surface fish for comparing population-specific genetic differences contributing to trait evolution. Nature Communications, 12(1), 1447.

Wilkens, H. (1988). Evolution and Genetics of Epigean and Cave Astyanax fasciatus (Characidae, Pisces). In M. K. Hecht and B. Wallace (Eds.), Evolutionary Biology: Volume 23 (pp. 271–367). Boston, MA: Springer US.

Xiong, S., Wang, W., Kenzior, A., Olsen, L., Krishnan, J., Persons, J., … Rohner, N. (2022). Enhanced lipogenesis through Pparγ helps cavefish adapt to food scarcity. Current Biology: CB, 32(10), 2272–2280.e6.

Xu, X., Guo, H., Zhang, Z., Wang, Y., Qin, J., and Zhang, X. (2021). Impact of pre- aggressive experience on behavior and physiology of black rockfish (Sebastes schlegelii). Aquaculture (Amsterdam, Netherlands), 536(736416), 736416.

Yoshizawa, M., Robinson, B. G., Duboué, E. R., Masek, P., Jaggard, J. B., O’Quin, K. E., … Keene, A. C. (2015). Distinct genetic architecture underlies the emergence of sleep loss and prey-seeking behavior in the Mexican cavefish. BMC Biology, 13(1), 15.

Yoshizawa, M., Yamamoto, Y., O’Quin, K. E., and Jeffery, W. R. (2012). Evolution of an adaptive behavior and its sensory receptors promotes eye regression in blind cavefish. BMC Biology, 10, 108.

Zabegalov, K. N., Kolesnikova, T. O., Khatsko, S. L., Volgin, A. D., Yakovlev, O. A., Amstislavskaya, T. G., … Kalueff, A. V. (2019). Understanding zebrafish aggressive behavior. Behavioural Processes, 158, 200–210.

