## Supplemental Figures for "Automated profiling of social behaviors to assess the genetic basis of evolution of aggressive behaviors in *A. mexicanus*"

**NODE PLACEMENT**

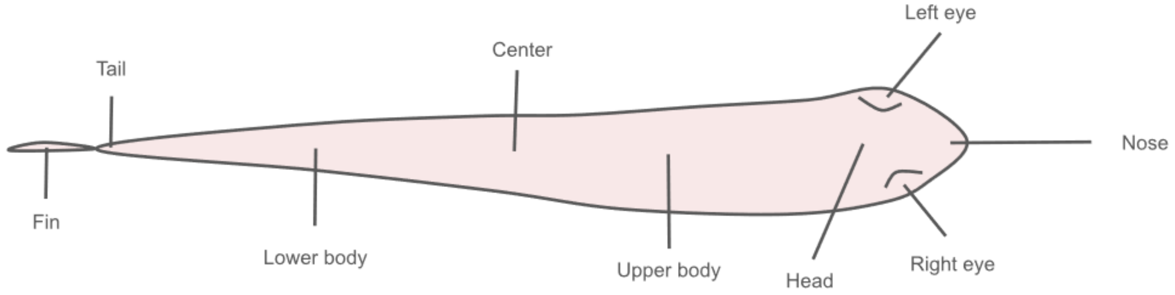


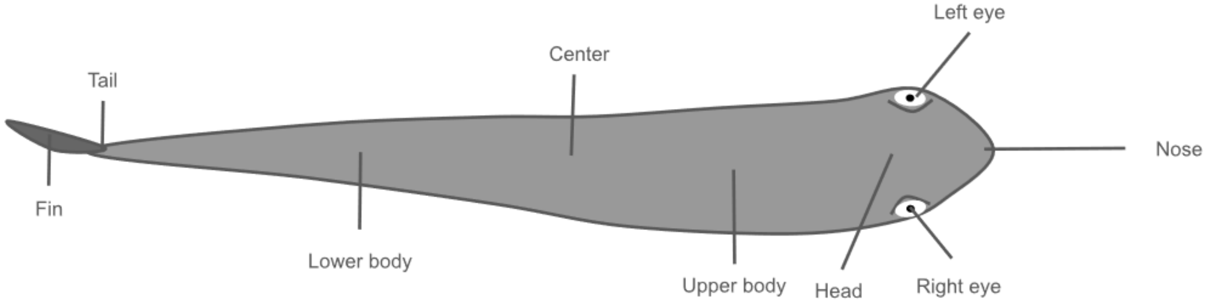


**AUTOMATED TRACKING USING SLEAP**

**AUTOMATED BEHAVIORAL ANNOTATION USING SIMBA**

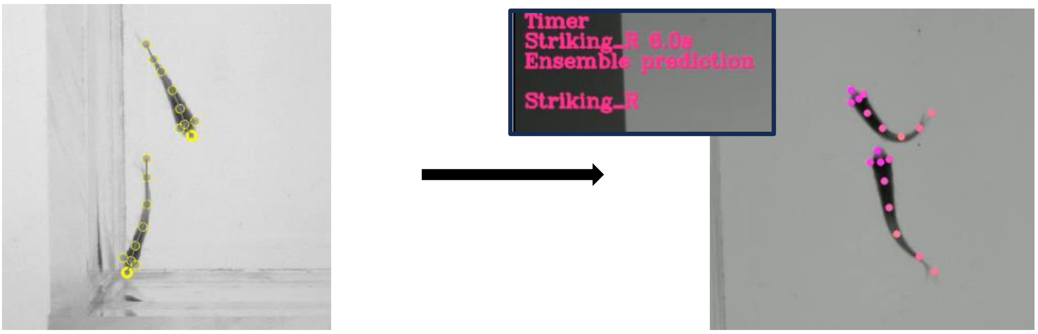


**Supplemental Figure 1.** Diagram depicting SLEAP node placement on Pachón and surface fish body parts. Still image depicting automated tracking using SLEAP showing the 9 nodes on each fish: nose, head, right eye, left eye, upper body, center, lower body, tail, and fin. Still image depicting automated behavioral annotation using SimBA showing the imported SLEAP tracks during a frame in which striking is predicted by the trained model. Images have been cropped to more easily visualize fish.

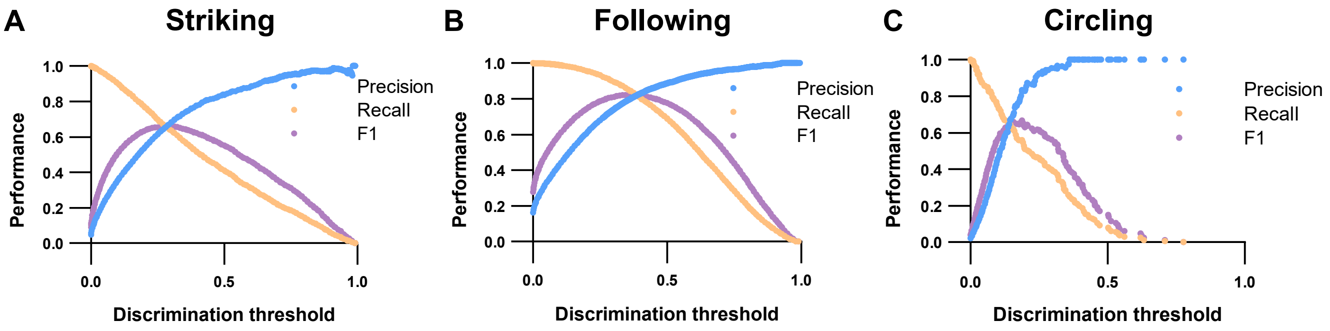


**Supplemental Figure 2.** Parameter optimization. Precision-recall-F1 curves for (A) striking, (B) following, and (C) circling behavioral classifiers to determine model performance and the optimal discrimination threshold parameters. The precision curve evaluates how many of the behavioral instance positives identified by the model are true positives. The recall curve evaluates how many of the total true positive instances were missed by the model. The F1 curve is the harmonic mean between the precision and recall curves.

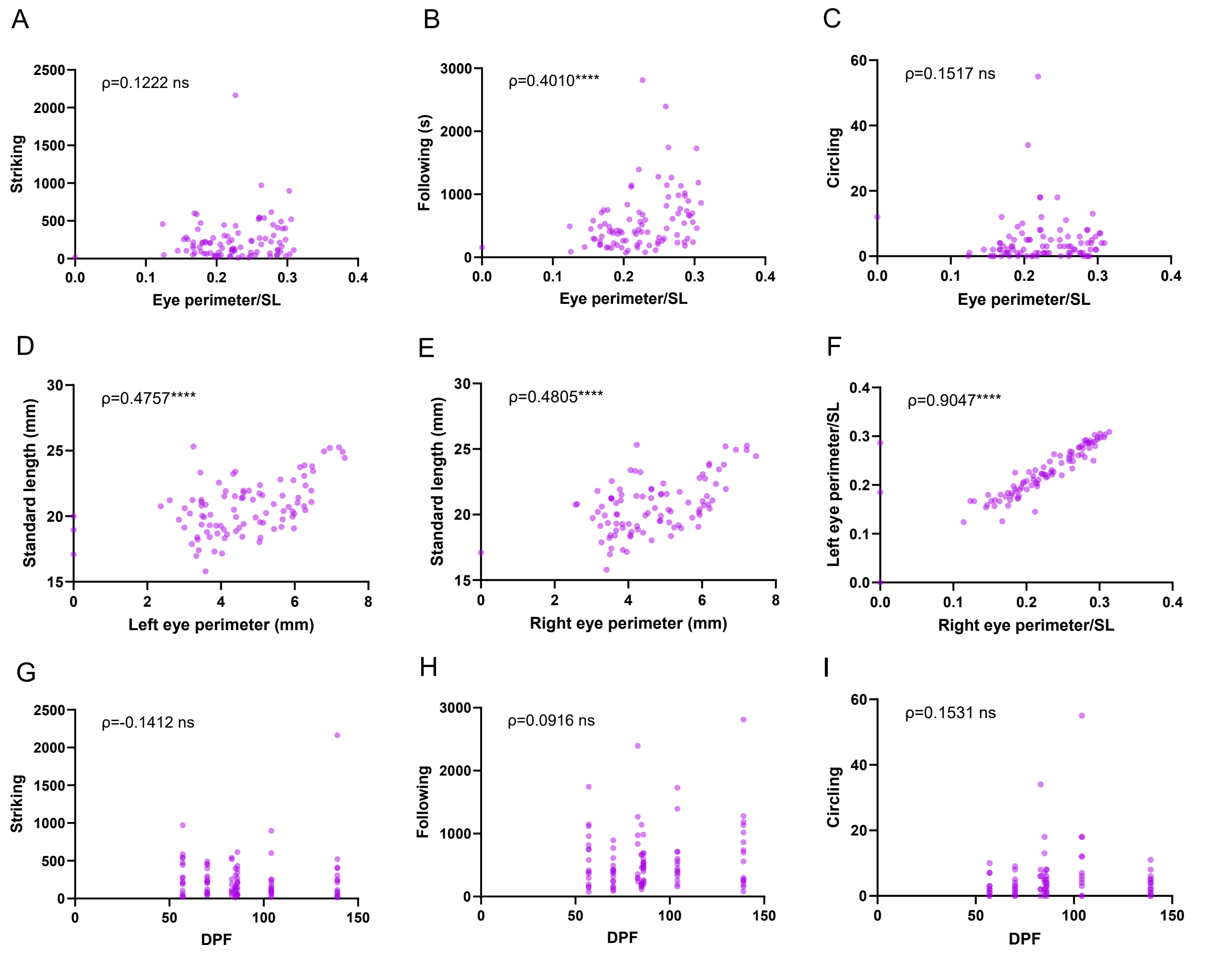


**Supplemental Figure 3.** Spearman correlations in surface-Pachon F2 hybrids. Correlations between relative left eye perimeter and (A) striking bouts (*rho*=0.1222, *p*=0.2407), (B) time following (s) (*rho*=0.4010, *p*<0.0001), and (C) circling bouts (*rho*=0.1517, *p*=0.1443) per hour. Correlations between standard length (mm) and (D) left eye perimeter (mm) (*rho*=0.4757, *p*<0.0001) and (E) right eye perimeter (mm) (*rho*=0.4805, *p*<0.0001). (F) Correlation between relative right eye perimeter and relative left eye perimeter (*rho*=0.9047, *p*<0.0001). Spearman correlations between striking and age recorded as days post fertilization (dpf) in (G) striking (*rho*=-0.1412, *p*=0.1677), (H) following (s) (*rho*=0.0916, *p*=0.3721), and (I) circling (*rho*=0.1531, *p*=0.1344).
